# Neural substrates and behavioral relevance of speech envelope tracking: evidence from post-stroke aphasia

**DOI:** 10.1101/2024.03.26.586859

**Authors:** Pieter De Clercq, Jill Kries, Jonas Vanthornhout, Robin Gerrits, Tom Francart, Maaike Vandermosten

## Abstract

Neural tracking of the low-frequency temporal envelope of speech has emerged as a prominent tool to investigate the neural mechanisms of natural speech processing in the brain. However, there is ongoing debate regarding the functional role of neural envelope tracking. In this context, our study aims to offer a novel perspective by investigating the critical brain areas and behavioral skills required for neural envelope tracking in aphasia, a language disorder characterized by impaired neural envelope tracking.

We analyzed an EEG dataset of 39 individuals with post-stroke aphasia suffering a left-hemispheric stroke who listened to natural speech. Our analysis involved lesion mapping, where left lesioned brain voxels served as binary features to predict neural envelope tracking measures. We also examined the behavioral correlates of receptive language, naming, and auditory processing (via rise time discrimination task) skills.

The lesion mapping analysis revealed that lesions in language areas, such as the middle temporal gyrus, supramarginal gyrus and angular gyrus, were associated with poorer neural envelope tracking. Additionally, neural tracking was related to auditory processing skills and language (receptive and naming) skills. However, the effects on language skills were less robust, possibly due to ceiling effects in the language scores.

Our findings highlight the importance of central brain areas implicated in language understanding, extending beyond the primary auditory cortex, and emphasize the role of intact auditory processing and language abilities in effectively processing the temporal envelope of speech. Collectively, these findings underscore the significance of neural envelope tracking beyond mere audibility and acoustic processes.

**Significance statement:** While some studies have proposed that neural envelope tracking primarily relates to audibility and acoustic speech processes, others have suggested its involvement in actual speech and language comprehension. By investigating the critical brain areas and behavioral skills essential in aphasia, we argue for a broader significance of neural envelope tracking in language processing. Furthermore, our findings highlight a specificity among individuals with aphasia, indicating its correlation with lesions in temporal brain regions associated with receptive language functions. This addresses the significant heterogeneity in lesion characteristics present among individuals with aphasia and suggests the potential of neural tracking as an EEG-based tool for specifically assessing receptive language abilities in this population.

## 1 Introduction

When processing speech, our brain exhibits a time-locked response to the stimulus. This response can be quantified by establishing the statistical relationship between recorded brain signals and speech representations of the stimulus. The strength of this statistical relationship is referred to as neural tracking (for a review, see (Brodbeck and Simon, 2020)). Neural tracking of various speech representations has been measured, but the temporal envelope of speech stands out as the most extensively studied speech representation in the literature (Gillis et al., 2022a). It is a pivotal cue for speech understanding (Shannon et al., 1995), encompassing cues for detecting and processing linguistic units of speech (e.g., phonemes, syllables and words) (Giraud and Poeppel, 2012). However, considerable debate exists in the literature about the functional role underlying neural envelope tracking. Some studies allocate a role in mere audibility and acoustic processes, and suggest speech understanding should be measured using higher-order linguistic speech representations (Verschueren et al., 2022; Gillis et al., 2023; Karunathilake et al., 2023; Kösem et al., 2023). Yet, other studies argue a role for neural envelope tracking in speech and language understanding, e.g. by demonstrating reduced tracking for incomprehensible speech or syntactically incorrect stimuli (Keitel et al., 2018; Etard and Reichenbach, 2019; Kaufeld et al., 2020; Coopmans et al., 2022).

Studies applying neural envelope tracking in language disordered populations, where linguistic deficits underlie the observed language problems, could provide a different perspective on this debate. Among various disorders, neural tracking has found application in the study of aphasia, a language disorder commonly caused by a stroke in the language-dominant left hemisphere of the brain (Papathanasiou and Coppens, 2017). In response to narratives, individuals with post-stroke aphasia (IWA) display decreased neural tracking of the envelope and linguistic speech representations (Kries et al., 2023a; De Clercq et al., 2023a, 2024b). Neural tracking measures further predicted aphasia at the individual level (De Clercq et al., 2023b), with prediction accuracy almost entirely driven by envelope tracking rather than linguistic tracking (De Clercq et al., 2024b). These findings may suggest a role for envelope tracking in language-related processes, given its ability to detect a language disorder through reduced envelope tracking measures.

However, the factors driving impaired neural envelope tracking of speech in aphasia remain elusive. Although linguistic abilities are assumed to underlie language deficits, IWA often present with lower-level auditory processing deficits as well, e.g., impaired rise time processing (Kries et al., 2023b). Hence, it is possible that impaired neural envelope tracking in aphasia did not reflect a core language deficit, but rather impaired auditory processing. Furthermore, IWA in prior studies exhibited large heterogeneity in brain lesions within the left hemisphere. Hence, the critically damaged brain areas associated with decreased neural envelope tracking are unknown at present. If neural envelope tracking is implicated in mere audibility and acoustic processes, it is expected that impaired neural envelope tracking associates with primary auditory cortex damage. Studies involving young listeners, however, have demonstrated that the envelope of speech sounds is not only encoded in the primary auditory cortex but extends to language-related areas such as the superior and middle temporal gyri (STG, MTG) and inferior frontal gyrus (IFG) (Kubanek et al., 2013; ten Oever et al., 2022).

In the present study, we investigate the correlates of impaired EEG-based neural envelope tracking in post-stroke aphasia, aiming to obtain a better understanding of the mechanisms driving neural envelope tracking. Specifically, we map neural envelope tracking measures to lesioned brain areas, a technique known as lesion mapping. This technique enables us to infer brain regions essential for maintaining intact neural envelope tracking. Furthermore, we investigate the link with performance on behavioral tests that asses core language skills, such as receptive language and naming, as well as lower-level auditory processing skills like rise time processing. Collectively, these analyses provide a better understanding of the mechanisms underlying neural envelope tracking and its clinical implications in aphasia.

## 2 Materials and methods

### 2.1 Participants

The presented sample is a subset of prior studies (Kries et al., 2023a; Mehraram et al., 2024), comprising 39 native Flemish IWA (13 female participants, mean age = 69.6 y/o, sd = 12.4 y/o) in the chronic phase (≥ 6 months) post-stroke. Compared to these prior studies, one participant was excluded as we lacked accessible brain scan data. Additionally, data from healthy age-matched controls described in previous studies were not included in the main analyses, as our focus was on investigating the link between brain lesions and behavioral impairments specific to the aphasia group. We recruited IWA at the stroke unit of the University Hospital Leuven and via speech-language pathologists. All IWA suffered a left-hemispheric or bilateral stroke, were diagnosed with aphasia in the acute stage after stroke using behavioral language tests and had no psychiatric or neurodegenerative disorder. Participant-specific demographic and lesion information is provided in the Extended Data, Table 1-1. For more information regarding recruitment strategy and diagnosis in the acute stage after stroke, we refer to Kries et al. (2023b). The study was approved by the ethical committee UZ/KU Leuven (S60007), and all participants gave written consent before participation. Research was conducted in accordance with the principles embodied in the Declaration of Helsinki and in accordance with local statutory requirements.

**Table 1.** Distribution of voxels that form the cluster surviving permutation testing across brain areas.

|  | Distribution voxels |
| --- | --- |
| MTG | 27.03% |
| SMG | 22.73% |
| Angular gyrus | 22.65% |
| Lateral occipital cortex | 15.71% |
| STG | 6.38% |
| Planum temporale | 2.31% |
| Planum polare | 2.39% |
| Heschl's gyrus | 0.73% |
| Insula | 0.07% |

### 2.2 Lesion segmentation and image processing

Lesion masks were segmented on T2-weighted FLAIR images for 37 participants, and on non-enhanced contrast computed tomography images for two participants. The lesions were manually segmented using MRIcron (v. 02092019), guided by the information documented in their respective medical files (written by a neurologist/neuroradiologist). Brain scans were acquired in the acute stage post-stroke (between 3-7 days) for 26 IWA, and 13 IWA had lesion data in post-stroke stage ≥ 4 months when participating in other studies performed in our research lab (Schevenels et al., 2022; De Clercq et al., 2024a). After segmentation, all brain scans and lesion masks were normalized to the template for older individuals in MNI space with the Clinical Toolbox implemented in SPM (Rorden et al., 2012). Finally, lesion masks were smoothed using a Gaussian kernel with a size of 4 mm using SPM12 (Welcome Trust Centre for Neuroimaging, London, UK), implemented in MATLAB 2021b (MathWorks, Massachusetts, USA). We used the Harvard-Oxford atlas (Kennedy et al., 1998) to define brain regions.

### 2.3 Behavioral tests

#### Behavioral measures of interest

We describe the behavioral tests of interest below. In previous studies at our research lab, IWA performed worse on these tests compared to healthy controls (Kries et al., 2023a,b; Mehraram et al., 2024).

#### Rise time test

The rise time discrimination task measures how well the rate of change in amplitude at the onset of a sound is processed. The task was presented as a 3-alternative forced choice task, where participants had to discriminate a deviant stimulus from two identical standard stimuli (Van Hirtum et al., 2019). The stimuli consisted of one-octave noise bands centered at 1 kHz and differed in their rise times. The reference stimuli had a rise time of 15 ms. The deviant (target) stimuli were computed to have rise times that decreased logarithmically in 50 steps from 699 until 15 ms. The task followed a one-up/two-down staircase procedure with target performance at 70.7%. The task ended once 8 reversals (i.e., changes in direction) were reached, or after a maximum of 87 trials. As this task targets the threshold at which differences in rise time can be detected by the participant, higher scores (i.e., higher thresholds) imply worse performance. For more information on this task and visualization of the procedure, we refer to Kries et al. (2023a). The task was presented using the software APEX (Francart et al., 2008) and stimuli were presented to the left ear at 70 dB SPL. Of all 39 participants, 31 succesfully completed this task, i.e., a certain rise time was reached. The remaining 8 participants did not complete the task as they experienced it as too difficult. These participants were removed from the analysis.

#### ScreeLing test

We administered the ScreeLing, a comprehensive aphasia test battery (Visch-Brink et al., 2010; El Hachioui et al., 2017). The ScreeLing is a validated aphasia battery testing for phonological, semantic and syntactic skills, primarily consisting of receptive test items. Each component contains four tasks with 24 items. In total, the maximum score on this test is 72, with lower scores reflecting worse performance. We used the total score to investigate associations between neural envelope tracking and general language skills in aphasia. One participant did not complete the ScreeLing test due to lack of time and was excluded from the analysis.

#### Naming test

We administered a separate test to asses naming abilities in aphasia, namely the Dutch picture-naming test (Nederlandse Benoem Test) (Van Ewijk et al., 2020), as the ScreeLing test does not include a naming subtest. This is a validated picture-naming test with 92 items and a maximum score of 276 points, with lower scores reflecting worse performance. One participant did not complete the naming test and was excluded from the analysis.

#### Behavioral measures that serve as covariates

We administered tests of hearing ability and cognition which serve as covariates. This was done as hearing abilities can impact neural envelope tracking outcomes (Gillis et al., 2022b). Furthermore, cognitive problems are often present in aphasia as well (El Hachioui et al., 2014; Rohde et al., 2018).

#### Hearing test

Hearing thresholds were assessed using pure tone audiometry (PTA) at octave frequencies ranging from .25 to 4 kHz. In case the hearing thresholds below 4 kHz were ≥ 25 dB HL, we augmented the stimulus presentation during the EEG experiment (standard at 60 dBA) with half the threshold at .25, .5 and 1 kHz per ear individually. This procedure was adapted from Jansen et al. (2012).

We also calculated the Fletcher index as the average of thresholds at 1, 2 and 4 kHz per ear. Subsequently, we calculated the average Fletcher Index across both ears and used this measure as a covariate in our statistical analyses.

#### Cognitive test

We administered the Oxford Cognitive Screen-NL (OCS) (Huygelier et al., 2019) to assess cognitive functions in our sample. The OCS is a validated test that measures cognition independent from language, hence allowing to disentangling cognitive from language functioning. We administered the subscales attention (i.e., crossing out target shapes among distractor shapes), executive functions (i.e., connecting circles and triangles in alternation in descending order of size) and memory (i.e., free recall and recognition of words and shapes from the attention subscale), and calculated a composite score of cognition, identical to the procedure in Kries et al. (2023b).

### 2.4 EEG experiment

We recorded participants’ EEG in a soundproof, electromagnetically shielded booth using a 64-channel BioSemi ActiveTwo system (Amsterdam, the Netherlands) at a sampling frequency of 8,192 Hz. During the EEG recording, participants were instructed to attentively listen to a story of approximately 25 minutes, *De Wilde Zwanen*, written by Christian Andersen and narrated by a female Flemish-native speaker. The story was cut into five parts with each an average duration of 4.84 minutes. After each story part, participants answered non-validated content questions about the preceding part, introduced to make the participant follow the content attentively. The story was bilaterally presented through ER-3A insert earphones (Etymotic Research Inc, IL, USA) at 60 dBA (but augmented in case of hearing loss with half the PTA threshold at .25, .5 and 1 kHz per ear individually) using the software platform APEX (Francart et al., 2008)

### 2.5 Signal processing

#### Envelope extraction

The envelope of the speech signal was extracted using a gammatone filter bank (Søndergaard et al., 2012) using 28 channels spaced by one equivalent rectangular bandwidth and center frequencies from 50 Hz until 5000 Hz. We extracted the envelopes of all 28 sub-bands by taking the absolute value of each sample and raising it to the power of 0.6 (Biesmans et al., 2015; Stevens, 1955). The envelopes of all sub-bands were then averaged to obtain a single envelope. The envelope was then downsampled using an anti-aliasing filter to 512 Hz to decrease processing time and filtered 0.5-40 Hz using a Least Squares filter of order 2000. We compensated for the group delay and used a transition band of 10% below the highand 10% above the lowpass frequency (i.e., highpass stopband at 0.45 Hz and lowpass stopband at 44 Hz). After filtering, the envelope was Z-score normalized and downsampled using an anti-aliasing filter to 128 Hz.

#### EEG data processing

EEG data were pre-processed using the Automagic toolbox (Pedroni et al., 2019) and custom MATLAB scripts (The MathWorks Inc., Natick, MA, USA, 2021). We first downsampled the EEG signals to 512 Hz to decrease processing time, and artifacts were removed using the artifact subspace reconstruction method (Mullen et al., 2015). Subsequently, we applied an independent component analysis, and components classified as “brain” or “other” (i.e., mixed components) with a probability higher than 50%, using the EEGLAB plugin ICLabel (Pion-Tonachini et al., 2019), were preserved (average of removed components=27, std=6; average preserved components=37, std:6). The neural signals were then projected back to the channel space, where the signals were common-average referenced and filtered in the same frequency band using the same Least Squares filter as in the envelope extraction method. Next, EEG signals were Z-score normalized and downsampled to 128 Hz.

### 2.6 Neural envelope tracking

We investigated neural envelope tracking using mutual information (MI) analysis, which is a nonlinear technique that outperforms common linear models and allows for spatial and temporal interpretations of the results (De Clercq et al., 2023b). We used the multivariate MI analysis, which determines the statistical relationship between the envelope and all 64 EEG channels combined. The MI analysis follows the Gaussian copula procedure as described in (Ince et al., 2017). In short, the envelope and the EEG data are separately ranked on a scale from 0 to 1, obtaining the cumulative density function (CDF). By computing the inverse standard normal CDF for each variable, the data distributions are transformed to perfect standard Gaussians. Subsequently, the parametric Gaussian MI estimate can be applied to the data provided by:

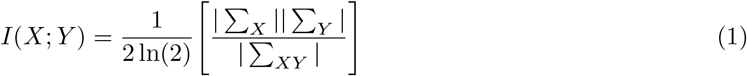

where I(X;Y) equals the MI between X and Y (i.e., the EEG and the envelope), expressed in bits. | ∑_*X*_ | and | ∑_*Y*_ | are the determinants of the covariance matrices of *X* and *Y*, and |∑_*XY*_ | is the determinant of the covariance matrix for the joint variable. To obtain temporal information on MI (i.e., the evolution of how the brain processes the speech stimulus over time), we compensated the delay between speech and the brain response by shifting the EEG relative to the envelope using an integration window of 0-500 ms, capturing brain latencies of interest. We calculated the mean MI over this integration window, obtaining a single outcome reflecting neural envelope tracking per participant. We refer to Ince et al. (2017) for an in-depth explanation of the Gaussian copula MI method. For a more practical explanation in the context of neural envelope tracking, see De Clercq et al. (2023b).

### 2.7 Statistical analysis

#### Link to lesion

To evaluate which lesioned brain areas are associated with neural envelope tracking, we conducted a univariate lesion mapping procedure, a common technique adapted from voxel-wise lesion-symptom mapping (Bates et al., 2003; Schwartz et al., 2009). In this approach, each voxel is used as a binary predictor (X, 1= lesioned voxel, 0= intact voxel) to predict the neural tracking outcomes (Y) using linear models. We treated demographic and lesion information (i.e., age, time post-stroke and lesion size) as nuisance variables, where all variance explained by the nuisance variables are removed from the outcome measures. Subsequently, the linear models are fit to the data. The obtained t-values were transformed to standardized Z-values for interpretation. Negative Z-values indicate that lesioned voxels relate to decreased envelope tracking. We restricted lesion mapping analyses to voxels in which at least 4 participants (i.e., at least 10%) share a lesion. Consequently, only voxels in the left hemisphere were included in the analyses. Analyses were performed using the NiiStat software (Rorden et al., 2007; Stark et al., 2019) implemented in MATLAB.

First, we report the unthresholded Z-values, and a thresholded value at p<.05 one-tailed (i.e., Z<-1.645) for exploratory purposes. For the main analysis, we correct for multiple comparisons employing a nonparametric cluster-based permutation procedure as suggested (Winkler et al., 2014) and implemented (Stark et al., 2019; Wilson et al., 2010) in prior work. We used 5000 permutations with cluster thresholding at p<.01 one-tailed (i.e., Z<-2.33). Permutation involves randomly shuffling the neural envelope tracking measures across participants while keeping the lesion maps fixed. Then, we applied the following procedure (see Stark et al. (2019) as well): 1) compute voxel-wise statistics for all 5000 permutations, 2) zero all voxels with Z>-2.33 (i.e., p>.01), 3) measure the cluster size of the largest surviving cluster per permutation, 4) rank the max cluster size of all 5000 permutations and 5) select the 50th largest cluster size (i.e., p=0.01). Then, we zero all voxels in the actual data with Z>-2.33 and of cluster size smaller than the cluster size determined in step 5 of the permutation procedure.

#### Link to behavior

We further explored the relationship between neural envelope tracking and behavioral measures of interest while controlling for covariates. We both applied frequentist and Bayesian statistics using RStudio (version 4.3.1). We used neural envelope tracking measures (i.e., MI in bits) as dependent variables, and behavioral measures of interest as independent variables, namely 1) the rise time threshold, 2) the ScreeLing test score and 3) the naming test score. We controlled for the following covariates: mean Fletcher Index across both ears (hearing), the test score on the OCS (cognition), age of the participant, the time post-stroke and lesion volume. Our linear models were constructed using the following formulas:

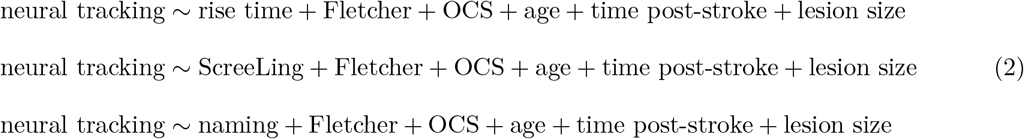

For frequentist statistics, we conducted a Type III ANOVA analysis and reported the F-statistic and p-value for each independent variable. P-values were adjusted for multiple comparisons using the false discovery rate (FDR) correction with the Benjamini-Hochberg procedure (Benjamini and Hochberg, 1995). For Bayesian statistics, we compared the full model as defined above to the null model, which includes only the covariates and not the independent variable of interest, using a uniform prior distribution. We report the Bayes Factor (BF10) of the full model compared to the reduced model, indicating the inclusion BF for the variable of interest. A BF10 value smaller than 1 indicates support for the null model (i.e., no evidence supporting inclusion of the variable), while a value larger than 1 reflects support for the alternative hypothesis.

For exploratory purposes, we also provide scatterplots and bivariate Spearman correlation coefficients (including BF10 calculation) between neural envelope tracking and behavioral measures of interest without correcting for covariates. P-values were also FDR-corrected for multiple comparisons. In the Extended Data, we also provide bivariate correlation coefficients between neural envelope tracking and covariates.

### 2.8 Data and code availability statement

All behavioral data and neural envelope tracking measures (pseudonymized), and code to reproduce the behavioral findings, will be made available upon publication. For the lesion mapping analysis, we used the NiiStat software package (Rorden et al., 2007; Stark et al., 2019) implemented in MATLAB. Our ethical approval does not permit public archiving of raw neuroimaging data, but raw EEG data can be made available upon request and if the GDPR-related conditions are met. The MRI data cannot be shared under any circumstance, as lesioned MRI data are person-specific and therefore cannot be considered anonymous.

## 3 Results

### 3.1 Lesion information

A lesion overlap map of our aphasia sample is provided in Figure 1. Largest lesion overlap was observed in the left insula and left inferior frontal gyrus (overlap of 19/39 participants). Although the majority of participants (23/39) had frontal lesions (inferior frontal gyrus, insula, precentral gyrus), many of these participants (12/23) had extensions of their lesion affecting temporal and parietal brain regions (Heschl’s gyrus, supramargynal gyrus (SMG), STG, MTG, temporal pole, angular gyrus, inferior temporal gyrus). In total, 23 out of 39 participants were faced with damage to some extent in the latter temporal and parietal brain regions. Additionally, 5 out of 39 participants only exhibited a lesion in other brain regions (occipital, cerebellum or midline), but were still classified as having aphasia as they failed behavioral diagnostic tests for aphasia and/or followed language therapy at the time of participation.

**Figure 1.**
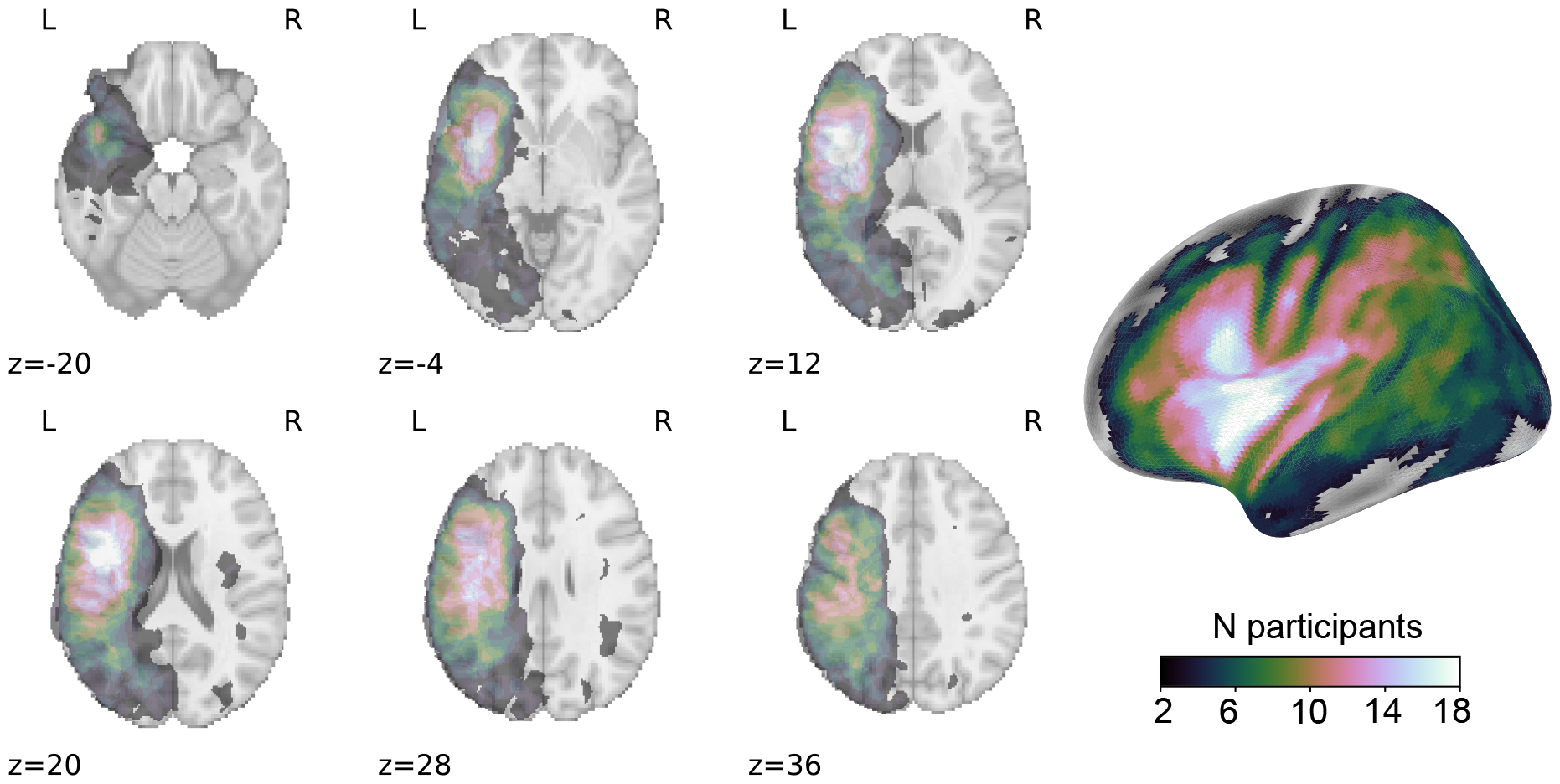
Lesion overlay image. Axial slices with corresponding MNI Z-coordinates are shown in neurological orientation. On the right, a surface image is shown.

### 3.2 Link to lesion

To explore the relationship between lesioned brain areas and neural envelope tracking, we employed a technique known as lesion mapping. In this approach, brain image voxels were used as binary predictors (1 = lesion, 0 = no lesion) to predict neural envelope tracking outcomes while controlling for covariates. Negative Z-values indicate that lesioned voxels are associated with poorer neural envelope tracking measures. The results are depicted in Figure 2, showing unthresholded Z-values (left) and Z-values after correction for multiple comparisons using cluster-based permutation testing (p<.01 one-tailed). A cluster encompassing temporal and parietal brain areas survived cluster-based permutation testing (a cluster size of 1254 voxels out of a total of 38196 voxels). The distribution of voxels in this cluster across different brain regions (using the Harvard-Oxford atlas) is provided in Table 1. The peak negative Z-value was observed in the MTG.

**Figure 2.**
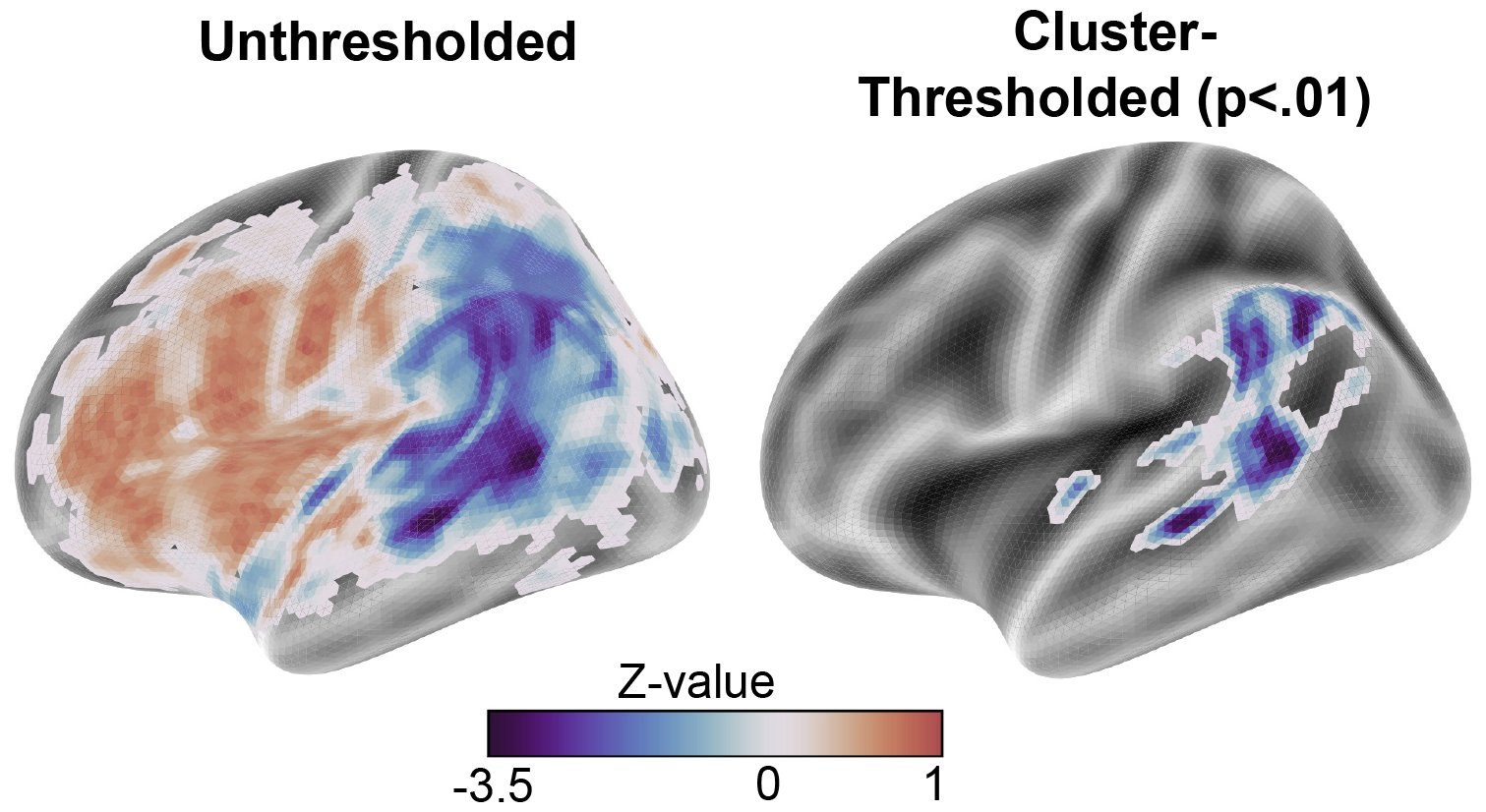
Lesion mapping results. Z-values represent the statistical relationship between lesioned voxels across subjects and neural envelope tracking outcomes.

### 3.3 Link to behavior

#### Spearman correlations

First, we present the bivariate Spearman correlations between neural envelope tracking and behavioral measures of interest. Scatterplots illustrating these correlations are provided in Figure 3. Neural envelope tracking showed significant correlations with the rise time threshold (Spearman R = -0.60, p < .001, BF10 = 68.77), the ScreeLing test score (R = 0.32, p = 0.047, BF10 = 2.71), and the naming test score (R = 0.35, p = 0.0435, BF10 = 1.57). Correlations and scatterplots between neural envelope tracking and behavioral (Fletcher, OCS score) and demographic (age, time post-stroke, lesion

**Figure 3.**
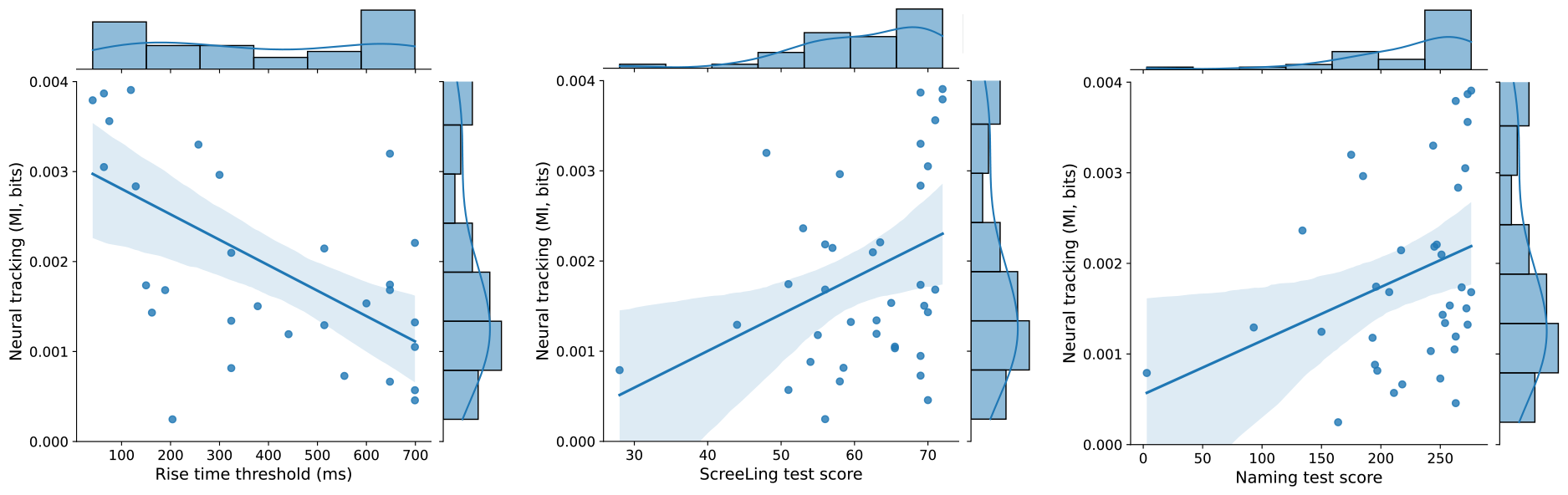
Scatterplots with regression lines. Visualization of the associations between neural envelope tracking and rise time threshold (left), SreeLing test score (middle) and naming test score (right). size) measures, which served as covariates, are provided in the Extended Data, Figure 1-1.

#### Linear models correcting for covariates

Finally, we examined the associations between neural envelope tracking and behavioral measures of interest while accounting for covariates using linear models as defined in Eq. 2. Type III ANOVA analyses revealed a main effect for the rise time threshold (F=8.96, p=0.018) but not for the ScreeLing test score (F=3.22, p=0.123) and the naming test score (F=2.12, p=0.156). We further calculated the inclusion BF for all behavioral scores, by comparing models as defined in Eq. 2 to the null model that only includes the covariates. The BF values all support the inclusion of the respective variable of interest over the null model (BF10=8.82 for the rise time threshold, BF10=1.74 for the ScreeLing test score, BF10=1.22 for the naming test score).

## 4 Discussion

The present paper investigated the lesion and behavioral correlates of neural envelope tracking in IWA. In terms of lesion analysis, our findings revealed that damage to brain areas adjacent to the primary auditory cortex, such as the MTG, angular gyrus, and SMG, was associated with diminished neural envelope tracking. This outcome implies a crucial role for language-related brain areas for neural envelope tracking of natural speech. Regarding behavioral observations, we found significant correlations and Bayesian support for associations between neural envelope tracking and both rise time processing and language skills. Taken together, these findings provide support for the involvement of language processes beyond mere audibility and acoustic processes in neural envelope tracking.

### 4.1 Lesion correlates of impaired neural envelope tracking in aphasia

The lesion mapping analysis revealed a cluster primarily comprising temporal and parietal brain areas, specifically the MTG, SMG and angular gyrus. These areas encompass core language-related brain areas known for their involvement in receptive language, phonological processing, and semantic processes (Hickok and Poeppel, 2007; Binder et al., 2009; Hartwigsen et al., 2010; Fedorenko et al., 2011; Price, 2012). The cluster also extended towards the lateral occipital cortex, which was somewhat unexpected. However, it is important to note that our analysis involved cluster identification, with peak Z-values and most voxels of the cluster observed in the MTG (see Table 1). Therefore, the MTG acted as a central hub driving the observed cluster. Additionally, spatial correlations within lesion maps (DeMarco and Turkeltaub, 2018) may explain the observed extensions towards occipital brain regions.

Our findings align with previous studies utilizing invasive recordings (Kubanek et al., 2013; Oganian and Chang, 2019) or MEG source reconstruction in healthy young listeners (Keitel et al., 2018; Brodbeck et al., 2018; ten Oever et al., 2022). For instance, Kubanek et al. (2013) showed that the envelope of nonspeech sounds is solely encoded in the primary auditory cortex, while speech sounds extend to language-related areas. ten Oever et al. (2022) demonstrated that neural envelope tracking of syntactically correct sentences exhibits stronger effects across the language network compared to tracking of syntactically incorrect stimuli or processing isolated syllables (ten Oever et al., 2022). These collective findings, combined with our novel outcomes at both the lesion and behavioral levels in a language-disordered population, suggest that higher-order language areas are critically involved in speech envelope processing.

Recent research has highlighted the impact of stroke on the conductivity of EEG signals (Piastra et al., 2022). This phenomenon can result in attenuated neural responses detected by EEG sensors, with more pronounced effects for lesions closer to the cortical surface. While this aspect requires further investigation, we maintain that these effects do not significantly alter our conclusions. Firstly, our findings indicate that lesions specifically affecting temporal brain areas are associated with reduced neural envelope tracking, despite the majority of IWA exhibiting frontal lesions (see Figure 1). Importantly, this analysis was adjusted for lesion size. Secondly, we did not observe an overall effect for brain areas close to the cortical surface throughout the left hemisphere. Instead, a clear division between temporal and frontal brain areas was evident. Finally, our observed behavioral correlates of impaired neural envelope tracking were also adjusted for lesion size, and we found no significant correlation between neural envelope tracking and lesion size (see Extended Data, Figure 1-1). Taken together, these results demonstrate specific location effects and behavioral consequences of impaired neural envelope tracking. Nonetheless, future studies should employ source reconstruction techniques as recommended by Piastra et al. (2022) and compare such neural tracking outcomes for IWA to those of healthy controls or a non-aphasic stroke group.

### 4.2 Behavioral correlates of impaired neural envelope tracking in aphasia

Our results indicated significant bivariate associations between neural envelope tracking and all behavioral test scores. However, effects were less robust for language test scores, reflected through non-significant effects after adjusting for covariates in an ANOVA analysis, and Bayesian factors were notably smaller compared to Bayesian factors for rise time processing. It is important to note that a ceiling effect on language test scores was observed in our sample, which was not present in the rise time thresholds (see distributions in Figure 3). This ceiling effect might have obscured a potential relationship with neural envelope tracking. Nonetheless, the lesion mapping analysis suggests an involvement of language functions, and Bayesian statistics still supported the alternative hypothesis of an effect of language tests on neural envelope tracking. Nevertheless, future studies should evaluate this relationship using more sensitive behavioral language measures or in more severely impaired aphasia samples. Alternatively, investigating the relationship with behavioral measures of produced, connected speech in aphasia (e.g., see (Le et al., 2018)) could provide insights into the same underlying speech processes as neural envelope tracking, particularly natural speech processes.

The functional role underlying neural envelope tracking is a subject of ongoing debate. Studies involving healthy young listeners typically compared different experimental conditions, such as clear speech to speech-in-noise (e.g., Vanthornhout et al. (2018)), syntactically correct sentences to incorrect sentences (e.g., Kaufeld et al. (2020)), or different languages (e.g., Gillis et al. (2023)). This paper provides a different perspective on the debate by studying which skills are critical for effective neural envelope tracking. Our results emphasize the significant role of rise time processing skills in this context. While our study cannot conclusively assign a role for neural envelope tracking in language skills at the behavioral level, trends in the data and a robust lesion effect in core language regions strongly support the involvement of neural tracking in higher-level language functions. Furthermore, hearing ability (as assessed with the Fletcher Index) was not associated with neural envelope tracking, and all analyses were corrected for this measure, arguing specificity of our measures beyond mere hearing skills in IWA.

Finally, our findings hold critical information for aphasia research. Envelope processing and auditory abilities in general have received limited attention in aphasia research over the past decades. Nevertheless, both the current paper and recent literature convincingly demonstrate the impairment of such fundamental auditory processes in aphasia at the behavioral (Kries et al., 2023b) and neural (De Clercq et al., 2023a; Kries et al., 2023a) level. Efficient rise time processing is, however, critical for higher-level speech comprehension as well, as these carry critical cues for detecting and segmenting phonemes, syllables and words (Goswami et al., 2011; Hämäläinen et al., 2012; Oganian and Chang, 2019). Furthermore, recent work has shown that impaired auditory processing skills can predict impaired phonetic processes in IWA, suggesting a cascading effect wherein deficits in lower-level auditory processing impact higher-order language functions (Kries et al., 2023b). Hence, we advocate for future research and clinical interventions to incorporate targeted plans addressing auditory processing impairments in aphasia. For example, a study implementing a training in auditory temporal processing improved language comprehension abilities in IWA (Szymaszek et al., 2017). Therefore, future studies should develop targeted interventions for auditory processing skills in aphasia and test their efficacy in language recovery.

### 4.3 Limitations and future directions

The lack of robustness in language effects prompts the need for future studies to employ more sensitive language measures (e.g., measures of produced connected speech, see Le et al. (2018)) or investigate severely impaired IWA. Furthermore, future studies should compare neural envelope tracking in IWA to stroke survivors without aphasia to further disentangle the effect of the lesion on the obtained outcome. Nevertheless, our findings already indicate the specificity of neural envelope tracking in lesion location across IWA. In addition, future studies could investigate the correlates of impaired higher-level linguistic speech tracking in aphasia. However, it’s crucial to note that these features also strongly correlate with the temporal envelope of speech (Gillis et al., 2022a), highlighting the necessity for complex statistical model comparisons involving different feature sets in a more extensive aphasia sample.

## 5 Conclusion

The current study delved into the neural substrates and behavioral correlates of diminished neural envelope tracking in post-stroke aphasia. Our results underscore the specificity of impaired neural envelope tracking within core language regions in the temporal and parietal cortex adjacent to the primary auditory cortex. At the behavioral level, Bayesian support was found for associations between neural envelope tracking and both auditory processing and higher-order language skills. These findings collectively point towards a role of neural envelope tracking in language functions beyond mere audibility and acoustic processes. Additionally, our findings deepen our comprehension of the clinical implications of impaired neural envelope tracking in aphasia, paving the way for EEG-based assessments targeting natural speech comprehension abilities in IWA.

## Acknowledgement

The authors would like to thank all participants. We would also like to extend our thanks to Dr. Klara Schevenels for her assistance in the recruitment process, as well as the individuals that helped with the data collection.

