## Supplementary Materials for "Neural substrates and behavioral relevance of speech envelope tracking: evidence from post-stroke aphasia"

### Extended Data

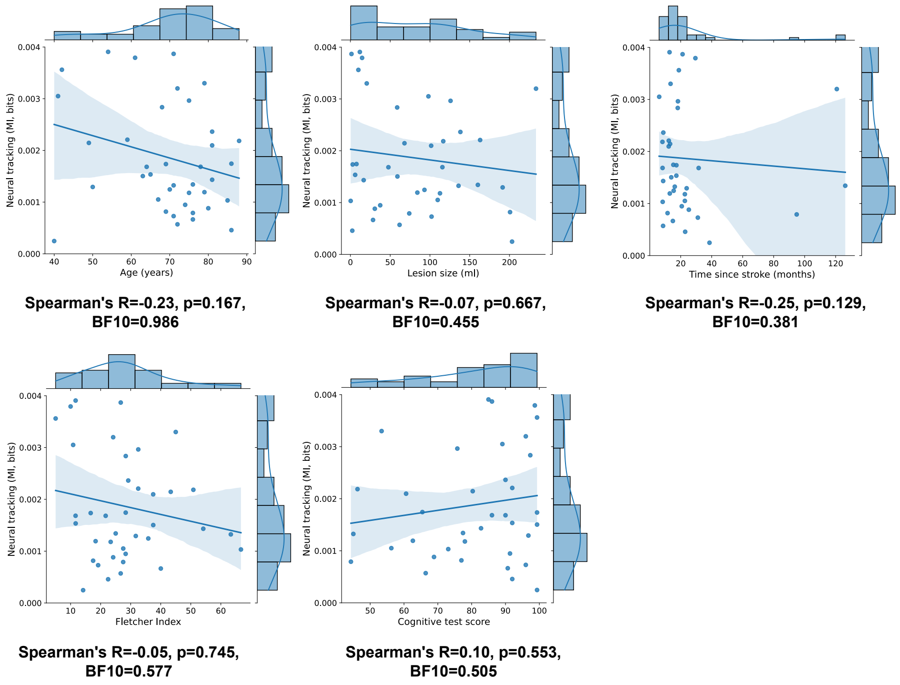


**Figure 1-1. Scatterplots between neural envelope tracking and covariates.** Statistics are provided for each figure (p-values not corrected for multiple comparisons).

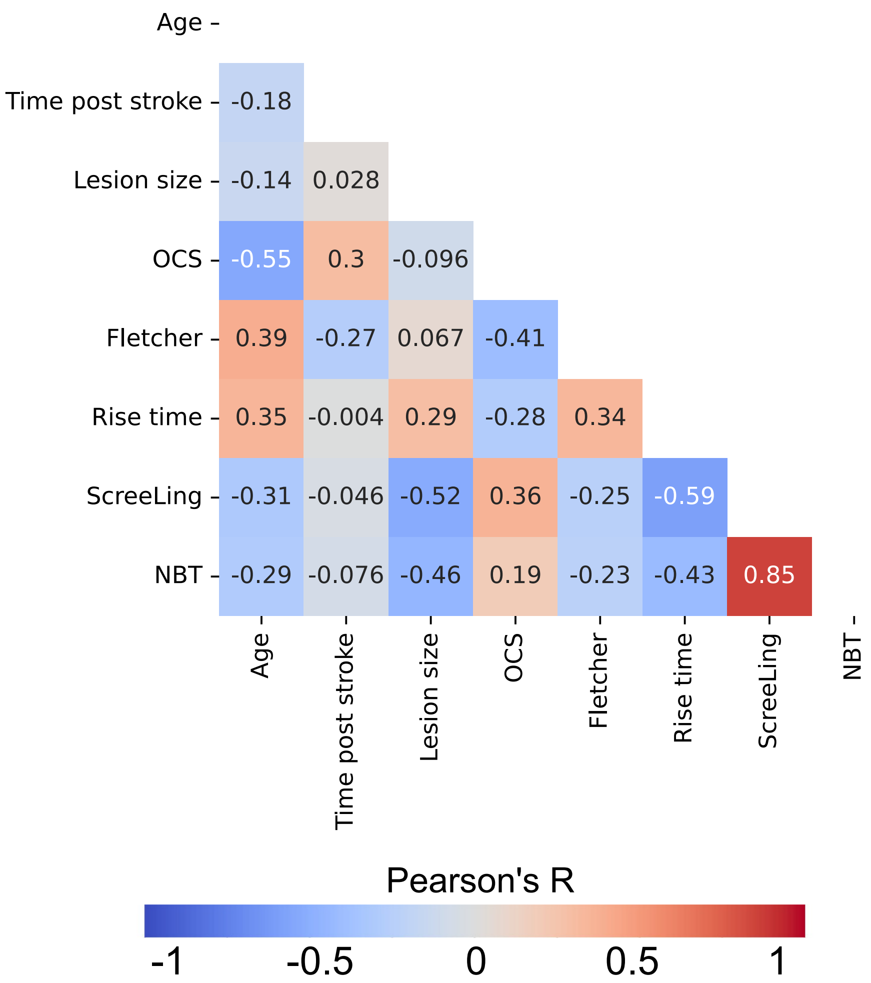


**Figure 1-2. Correlation matrix between all behavioral measures.** Spearman correlation coefficients among behavioral and demographic measures used in the main analyses.

**Table 1-1.** Demographics and lesion information

| ID | age | sex | Time since stroke (months) | Stroke type | Blood vessel | Lesioned hemi-sphere | Lesion size (ml) | SLT | Naming (max=276) | ScreeLing (max=72) |
|---|---|---|---|---|---|---|---|---|---|---|
| sub-006 | 86 | m | 22.8 | ischemia | VA | bilateral | 2.36 | yes | 263 | 70 |
| sub-008 | 71 | f | 21.1 | ischemia | MCA | left | 1.35 | no | 273 | 69 |
| sub-009 | 67 | m | 22.6 | ischemia | MCA | left | 109.73 | no | 262 | 65.5 |
| sub-010 | 74 | m | 20.6 | ischemia | PCA | left | 37.46 | no | ◇ | 69 |
| sub-014 | 75 | m | 18.2 | ischemia | MCA | left | 125.89 | yes | 185 | 58 |
| sub-016 | 68 | m | 18.0 | ischemia | MCA | bilateral | 58.78 | yes | 265 | 69 |
| sub-017 | 88 | m | 8.2 | ischemia | MCA/ PCA | bilateral | 116.73 | no | 245 | 56 |
| sub-018 | 61 | f | 29.4 | ischemia | MCA | left | 14.68 | yes | 263 | 72 |
| sub-019 | 72 | m | 8.6 | ischemia | PCA | left | 61.79 | ◇ | 211 | 51 |
| sub-020 | 42 | m | 18.7 | ischemia | MCA | left | 10 | yes | 273 | 71 |
| sub-021 | 81 | m | 8.6 | hemorrhage | PCA | left | 16.67 | no | 252 | 70 |
| sub-022 | 78 | f | 8.2 | hemorrhage | PCA | left | 115.65 | yes | 207 | 56 |
| sub-024 | 69 | m | 6 | ischemia | MCA | left | 116.9 | yes | 260 | 68.5 |
| sub-025 | 69 | m | 11.5 | ischemia | MCA | left | 200.69 | yes | 197 | 58.5 |
| sub-026 | 71 | m | 31 | ischemia | MCA | left | 102.01 | yes | 250 | 69 |
| sub-027 | 76 | m | 126.2 | ischemia | MCA | left | 160.23 | yes | 254 | 63 |
| sub-028 | 80 | m | 25.2 | ischemia | PCA | left | 30.68 | no | 195 | 54 |
| sub-029 | 75 | m | 22.6 | ischemia | MCA | left | 112.61 | yes | 193 | 55 |
| sub-030 | 49 | m | 13.1 | hemorrhage | MCA | left | 67.97 | yes | 217 | 57 |
| sub-031 | 79 | m | 12.9 | ischemia | MCA | left | 84.16 | yes | 263 | 63 |
| sub-032 | 76 | m | 94.8 | ischemia | MCA | left | 73.97 | yes | 3 | 28 |
| sub-034 | 79 | m | 13.4 | ischemia | PCA | bilateral | 20.52 | yes | 244 | 69 |
| sub-035 | 64 | f | 31.5 | ischemia | MCA/ ACA | left | 47.77 | yes | 276 | 71 |
| sub-049 | 85 | f | 8.3 | ischemia | PICA | left | 0.33 | yes | 242 | 65.5 |
| sub-050 | 81 | f | 8.7 | ischemia | MCA | left | 138.26 | yes | 134 | 53 |
| sub-052 | 72 | m | 120.7 | ischemia | MCA | left | 233.33 | yes | 175 | 48 |
| sub-053 | 41 | f | 6.1 | ischemia | MCA | left | 98.08 | yes | 271 | 70 |
| sub-054 | 69 | m | 17.2 | ischemia | MCA | left | 2.83 | yes | 268 | 69 |
| sub-056 | 76 | m | 14.9 | ischemia | MCA | left | 28.44 | yes | 218 | 58 |
| sub-059 | 86 | m | 15.2 | ischemia | ICA | left | 7.63 | no | 196 | 51 |
| sub-060 | 70 | m | 15.4 | ischemia | MCA | left | 93.1 | yes | 150 | ◇ |
| sub-061 | 54 | f | 12.7 | ischemia | MCA | left | 11.85 | yes | 276 | 72 |
| sub-063 | 63 | f | 13.9 | ischemia | MCA | left | 59.1 | yes | 272 | 69.5 |
| sub-064 | 81 | f | 12.2 | ischemia | crypto. | bilateral | 101.52 | yes | 251 | 62.5 |
| sub-065 | 50 | f | 23.8 | ischemia | MCA | left | 191.38 | yes | 93 | 44 |
| sub-067 | 40 | f | 38.5 | ischemia | MCA | left | 203.22 | yes | 164 | 56 |
| sub-068 | 59 | m | 12.2 | ischemia | MCA | left | 163.08 | no | 247 | 63.5 |
| sub-069 | 65 | m | 17.1 | ischemia | MCA | left | 5.98 | no | 258 | 65 |
| sub-070 | 71 | f | 16.1 | ischemia | MCA | bilateral | 135.93 | no | 273 | 59.5 |
| N=39 | 70<br>± 12 | 26m<br>13f | 24<br>± 28 | 36 ischemia<br>3 hemorrhage | 27 MCA<br>7 PCA<br>1 ACA<br>1 VA<br>1 PICA<br>1 ICA<br>1 crypto. | 33 left<br>6 bilateral | 81.09<br>± 65.07 | 28 yes<br>10 no | 244.71<br>± 58.11 | 61.55<br>± 9.40 |

SLT=speech-language therapy; VA=vertebral artery; MCA=middle cerebral artery; PCA=posterior cerebral artery; ACA=anterior cerebral artery; PICA=posterior inferior cerebellar artery; ICA=internal carotid artery; crypto.=cryptogenic; ◇=data not available
